# Stress–recovery dynamics reveal a separatrix in transcriptomic landscapes across organs and aging

**DOI:** 10.64898/2026.09.29.755524

**Authors:** Bingxian Xu, Shuqin Ji, Ziyi Lin, Chunhua Deng, Lei Li

## Abstract

Aging is often characterized statically as molecular profiles, but how these profiles may affect the capacity of systems to respond to the external environment remains elusive. We perturbed young and aged mice via transient sleep deprivation and profiled transcriptomes from six organs immediately after stress and following recovery. The same perturbation induced complex temporal dynamics for each gene in an organ- and age-dependent manner. A shape parameter distinguished genes that recovered toward control expression from those that continued to drift away during recovery. Transcriptomic vector fields were inferred from these transitions using neural ordinary differential equations, revealing distinct stability regimes across organ systems. Interestingly, that of testis from the young mice contained a separatrix bounding the region within which perturbed state can return toward control. These findings establish transient perturbation and recovery as a framework for probing age-dependent transcriptomic stability and identifying boundaries that determine whether a perturbed system can return.

## INTRODUCTION

Aging progressively alters molecular and cellular programs across tissues, leading to loss of physiological function and increased vulnerability to disease^1,2^. Multi-tissue transcriptomic studies have revealed both shared and tissue-specific features of aging, establishing detailed molecular maps of differences between young and aged states^3–5^. These studies have predominantly relied on static comparisons. Although such comparisons identify age-associated molecular changes, they do not directly assess the stability of a biological state or its capacity to return toward baseline following perturbation.

The ability to withstand and recover from a stressor is a central feature of biological resilience and is thought to decline during aging^6,7^. From a dynamical-systems perspective, transient perturbation followed by unconstrained recovery can provide information about local stability that is inaccessible from static measurements alone. Gene-expression states may be represented conceptually as positions within a landscape^8,9^, where perturbations displace a system and its subsequent trajectory reflects the geometry of its surrounding landscape^10,11^. The transcriptomic state of biological systems changes with age, yet how aging can impact the stability of the system remains poorly understood.

Sleep deprivation provides a transient systemic challenge with molecular consequences extending beyond the brain to peripheral tissues^12–14^. Its removal also creates a defined recovery period during which transcriptional states can return toward baseline, remain persistently displaced, or continue toward alternative states. This paradigm therefore offers an opportunity to probe how age and organ identity influence both the initial stress response and the trajectory that follows.

Here, we subjected young and aged mice to 24-h sleep deprivation and profiled transcriptomes from the adrenal gland, heart, liver, testis, hypothalamus, and brainstem immediately after stress or following 3 days of recovery. We quantified organ- and age-specific responses, classified gene-level trajectories using a shape parameter, and applied neural ordinary differential equations^15^ to infer vector fields describing transcriptomic state transitions. The reconstructed landscapes revealed distinct recovery regimes across organs and ages. In the young testis, the inferred vector field contained a separatrix bounding the region from which transcriptomic states can return toward control. These findings establish transient stress and recovery as a framework for probing transcriptomic stability and identifying boundaries that constrain restoration of biological states during aging.

## RESULTS

### Stress–recovery profiling probes transcriptomic landscapes across organs and aging

Static comparisons between young and aged organisms identify age-associated molecular change but do not directly reveal what really matters: can the system respond to and recover from the next perturbation? To answer this, we used a transient challenge followed by an unperturbed recovery period to probe transcriptomic stability across organs and ages.

Mathematically, a system of ordinary differential equations (ODEs), 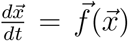, can characterize processes happening within an organism. In this expression, 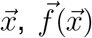 and *t* denote the quantity of species (mRNA, protein) involved, reaction kinetics and time respectively. We can assume that the system sits in some kind of steady state, 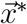, such that 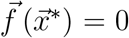. The stability of the state, 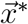, is probed by applying an infinitesimal perturbation 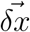. The effect of this perturbation decays to zero if the largest eigenvalue of the Jacobian matrix (*λ*_max_), **J**, of 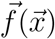 at 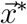 is smaller than 0 (Figure 1A). This “kinetic” view of the system can be transformed to a “landscape” view if we define another function 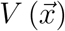 such that 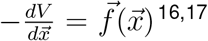. The “landscape” view offers an intuitive understanding of stability, where the system is stable if it is sitting at the bottom of the valley and unstable vice versa (Figure 1A). The mathematical definition of stability provided a key experimental insight: the system should be perturbed only transiently before allowing it to relax freely. In addition, one should not change the reaction kinetics, 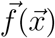, or equivalently the shape of the landscape, 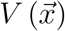 (Figure 1A). This framework provided the rationale for transiently perturbing the system and then allowing it to recover without continued intervention. The recovery period reveals whether the transcriptomic state returns toward its original position or remains displaced following removal of the challenge (Figures 1A and 1B).

**Figure 1.**
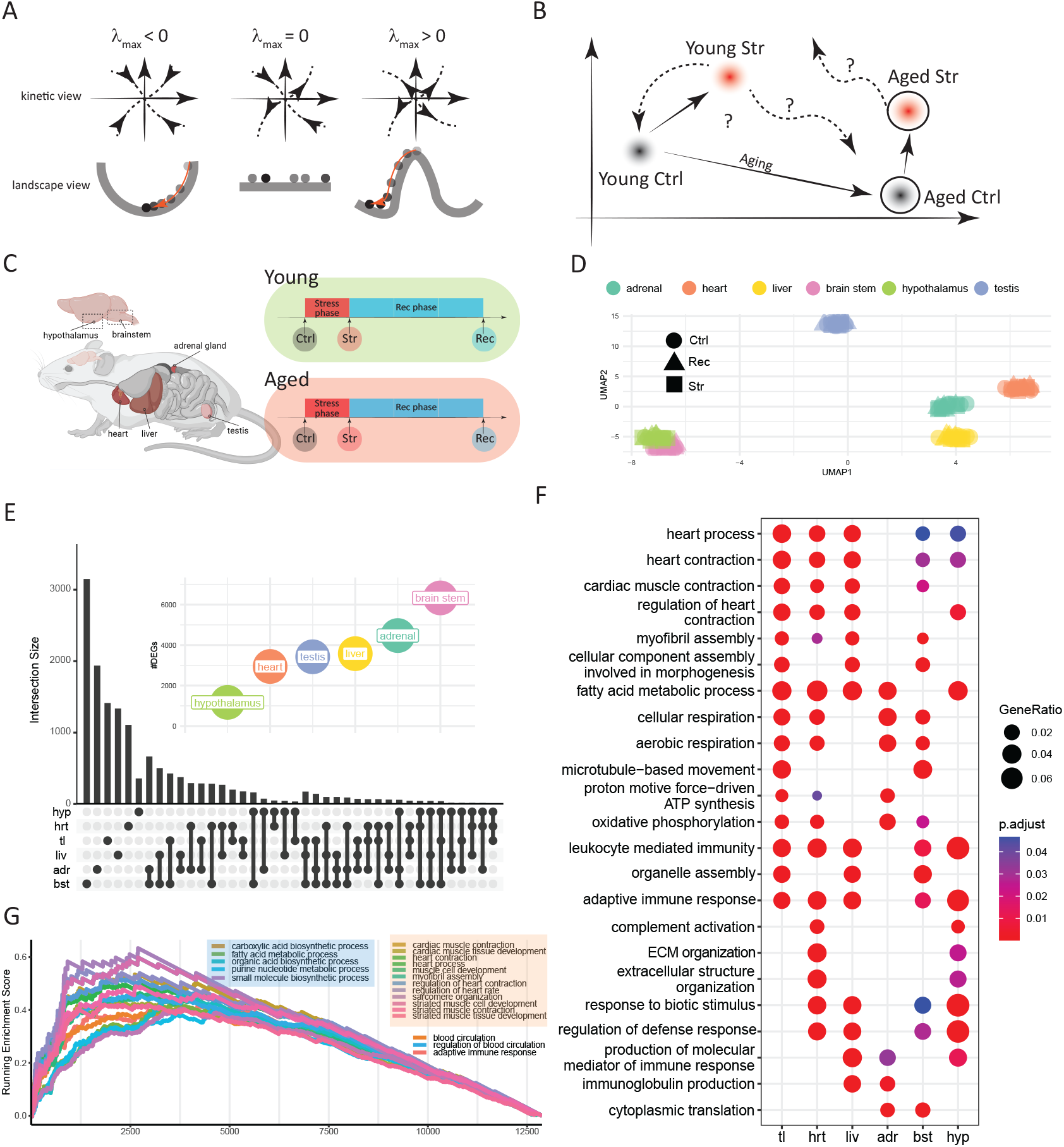
A stress–recovery framework for probing transcriptomic landscapes across organs and aging. (A)Relationship between the kinetic and the landscape view of gene expression models. A fixed point of a system is stable if its largest Jacobian eigenvalue is smaller than 0, unstable if its largest Jacobian eigenvalue is greater than 0 and a saddle if its largest Jacobian eigenvalue is equal to 0. A stable/unstable fixed point attracts/repels nearby points and is associated with a local landscape that is concaving up/down. (B)Illustrative trajectories that the system can take on the phase plane during aging and stress. (C)Experiment flowchart and sampling of organs (Ctrl, control; Str, sleep-deprivation stressed, Rec, sleep-deprivation stressed followed by 3-day recovery). (D)Visualization of the gene expression profiles on the UMAP space, colored by their organ of origin. (E)Upset plot of genes showing differential expression during aging in each organ. Intersecting DEGs across organs were indicated by the connected dots. Results revealed a highly organ-specific aging pattern (hyp, hypothalamus; hrt, heart; tl, testis; liv, liver; adr, adrenal gland; bst, brainstem). (F)Gene ontology terms enriched in each sampled organs when comparing the control groups of the young and aged cohorts. Enriched terms were related to muscle contraction, metabolism and immune response. (G)Enrichment plot from gene set enrichment analysis (GSEA) of common differentially expressed genes during aging in sampled organs. Result revealed common DEGs to be involved in metabolism, immune response and muscle contraction.

Young and aged male mice were assigned to untreated control (Ctrl), 24-h sleep deprivation (Str), or 24-h sleep deprivation followed by a 3-day recovery period in their home cages (Rec). The adrenal gland, heart, liver, testis, hypothalamus, and brainstem were collected for bulk RNA sequencing, generating transcriptomic measurements across organ, age, stress, and recovery conditions (Figure 1C). Uniform manifold approximation and projection (UMAP)^18,19^ separated samples primarily according to organ identity (Figure 1D). Samples from the same organ formed compact clusters, whereas age- and treatment-associated differences were not resolved at this level of visualization. Organ identity therefore accounted for the dominant global transcriptomic variation, indicating that aging, stress, and recovery should be evaluated within each organ.

We first compared untreated young and aged mice to establish age-associated transcriptomic differences. The brainstem exhibited the largest number of differentially expressed genes (DEGs), exceeding that detected in the hypothalamus by more than fivefold (Figure 1E). Despite extensive changes within individual organs, only five genes—*Trem2, Grn, Unc93b1, Ctla2a*, and *Gstm1*—were differentially expressed across all six organs. The limited intersection demonstrated that aging was associated predominantly with organ-specific changes at the individual-gene level. Despite this limited gene-level overlap, Gene Ontology over-representation analysis^20,21^ identified recurrent functional themes. Age-associated DEGs were enriched for processes related to metabolism, contractile functions, and immune regulation (Figure 1F). Consistently, gene set enrichment analysis^22^ of genes ranked by the number of organs in which they were differentially expressed identified related functional programs (Figure 1G). Thus, aging affected partially shared biological functions through largely organ-specific transcriptional changes.

### Stress and recovery engage organ- and age-specific transcriptional trajectories

We next investigated how age influenced the transcriptional response to the same stress challenge. Sleep deprivation induced more DEGs in the young adrenal gland, hypothalamus, and heart than in their aged counterparts, but fewer DEGs in the young brainstem, testis, and liver (Figure 2A). Aging therefore did not uniformly increase or decrease the breadth of the stress response; instead, its effect differed among organs.

**Figure 2.**
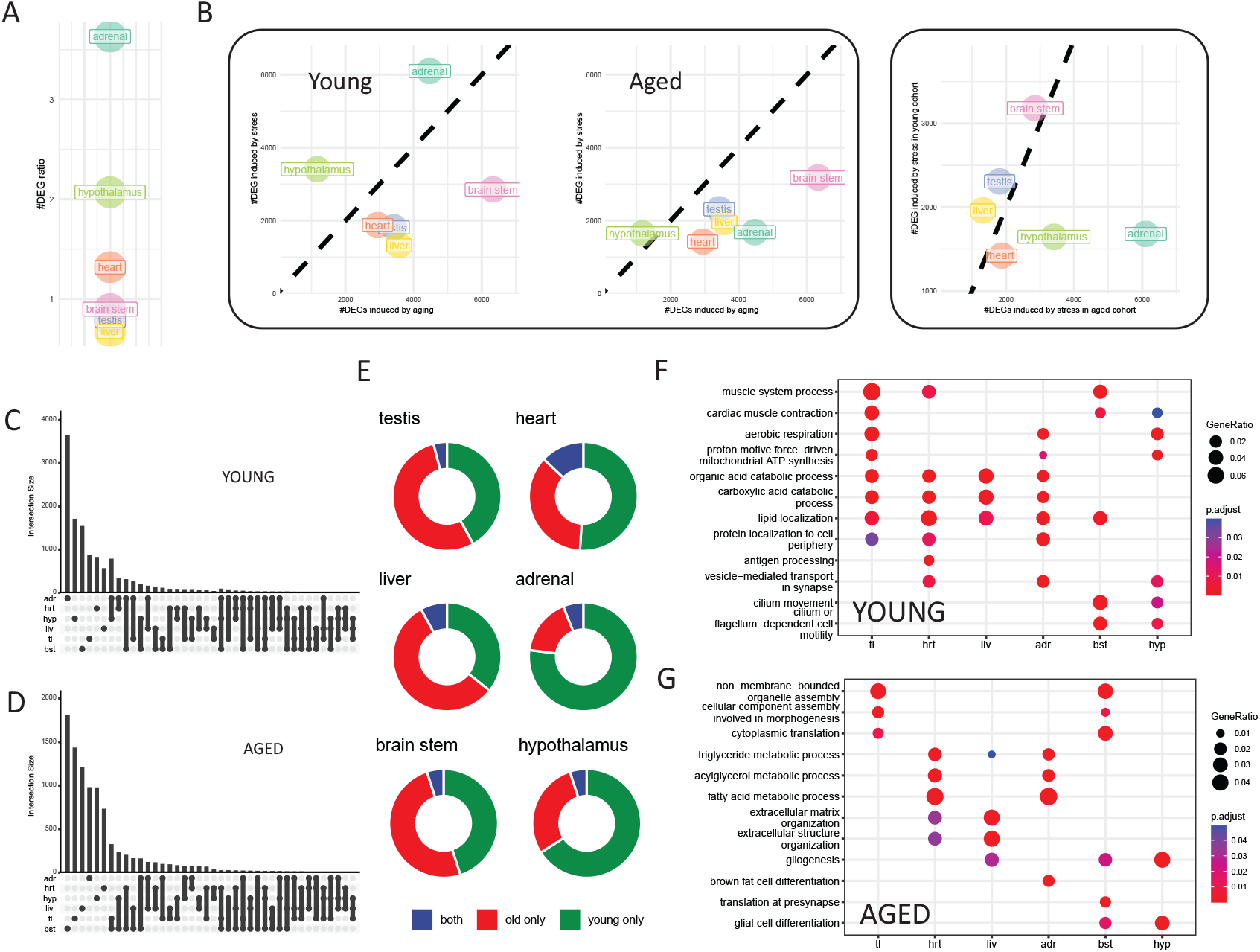
Organ- and age-specific transcriptomic responses to sleep deprivation stress. (A)Ratio of the number of sleep deprivation-responsive differentially expressed genes (DEGs) in young mice to that in aged mice for each organ. A ratio of 1 indicates equal numbers of responsive DEGs in the two age groups; a ratio greater than 1 indicates more responsive DEGs in young mice, whereas a ratio below 1 indicates more responsive DEGs in aged mice. (B)Relationship between the number of age-associated DEGs and the number of sleep deprivatio responsive DEGs across organs (left), and between the numbers of sleep deprivation-responsive DEGs in young and aged mice (right). (C)UpSet plot showing intersections of sleep deprivation-responsive DEGs across organs in young mice. (D)UpSet plot showing intersections of sleep deprivation-responsive DEGs across organs in aged mice. (E)Proportions of sleep deprivation-responsive DEGs detected only in young mice, only in aged mice, or in both age groups for each organ. (F)Gene Ontology over-representation analysis of sleep deprivation-responsive DEGs in young mice. (G)Gene Ontology over-representation analysis of sleep deprivation-responsive DEGs in aged mice. Sleep deprivation-responsive DEGs were identified by comparing the stress (Str) and untreated control (Ctrl) groups within each age and organ. Rec, sleep deprivation followed by 3-day recovery.

The number of age-associated DEGs was not significantly correlated with the number induced by sleep deprivation across organs (Figure 2B). Thus, the extent of baseline age-associated transcriptional change did not predict the number of genes recruited in response to the challenge. The response to sleep deprivation was highly organ-specific, with no single DEG shared across all six organs in either young or aged mice (Figures 2C and 2D). Only a small proportion of responsive genes in each organ were shared between the two age groups (Figure 2E), suggesting that the young and aged mice engaged largely distinct gene sets in response to the same perturbation.

Gene Ontology analysis revealed that these gene-level differences were accompanied by distinct functional responses. In young mice, sleep deprivation-responsive genes in the testis, heart, liver, and adrenal gland were enriched for metabolic processes, including lipid localization, organic acid catabolism, and steroid metabolism (Figure 2F). Genes associated with cilium movement were enriched in the brainstem and hypothalamus. This enrichment suggests the potential involvement of ciliated cells, such as ependymal cells lining the brain ventricles, whose significance in aging is well documented^23–25^. In aged mice, responsive genes in the testis and brain-stem were enriched for protein localization, cytoplasmic translation, and organelle assembly. The heart and liver exhibited partially overlapping responses involving triglyceride metabolism, immune-response activation, and extracellular-matrix organization (Figure 2G). Thus, both the genes and biological processes engaged by sleep deprivation varied across organs and ages.

We then examined how these responses changed following 3 days of recovery. We used the number of residual DEGs between the Rec and corresponding Ctrl groups as an operational measure of transcriptomic displacement after recovery. A reduction relative to the Str-versus-Ctrl comparison indicated movement toward the untreated control state but did not necessarily imply complete recovery. The hypothalamus was the only organ showing fewer residual DEGs after recovery in both age cohorts. Other organs showed signs of recovering in only one age group; for example, testis and heart only recovered in the aged and young cohort respectively (Figure 3A). Return toward the control state was therefore both organ- and age-dependent. The number of residual DEGs after recovery was weakly correlated with the number of age-associated DEGs across organs (*ρ* = 0.47, *p* = 0.13). This association indicated that organs with more extensive age-associated changes tended to retain more transcriptomic differences after recovery, but it did not establish a causal relationship between the rate of aging and recovery capacity.

**Figure 3.**
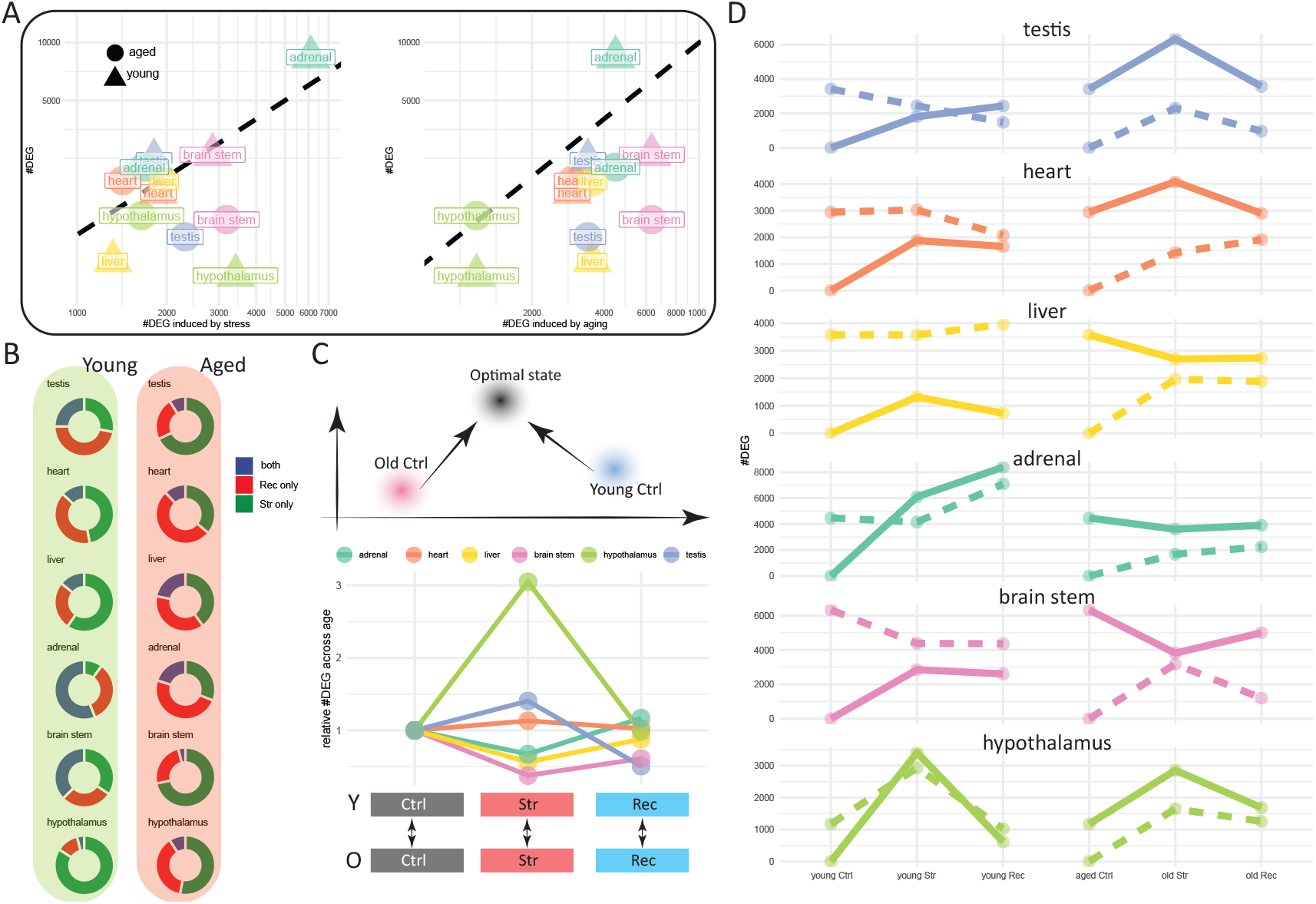
Organ- and age-specific transcriptomic trajectories during stress recovery. (A)Relationship between the number of residual differentially expressed genes (DEGs) after recovery (Rec versus Ctrl; y axis) and the number of DEGs induced by sleep deprivation (Str versus Ctrl; left), or associated with aging (Aged Ctrl versus Young Ctrl; right), across organs. The number of residual DEGs after recovery was weakly correlated with the number of age-associated DEGs. (B)Proportions of genes differentially expressed only immediately after sleep deprivation (Str-specific), only after recovery (Rec-specific), or under both conditions (persistent) in each organ and age group. DEGs were defined relative to the corresponding untreated control group. (C)Conceptual and transcriptomic assessment of convergence between young and aged states. Top, schematic illustrating the possibility that young and aged organs approach a shared transcriptomic state through different trajectories following the same perturbation. Bottom, transcriptomic difference between young and aged mice under the Str or Rec condition normalized to the difference between their corresponding Ctrl groups. A value below 1 indicates that the young and aged transcriptomic states are more similar under the indicated condition than under the Ctrl condition. (D)Changes in the numbers of DEGs relative to the Young Ctrl state (solid line) and Aged Ctrl state (dotted line) across the ordered Ctrl, Str, and Rec conditions. The two reference comparisons provide complementary measures of transcriptomic displacement and do not constitute a one-dimensional biological-age score. Ctrl, untreated control; Str, 24-h sleep deprivation; Rec, 24-h sleep deprivation followed by 3-day recovery.

We next classified genes according to whether they were differentially expressed only immediately after sleep deprivation, only after recovery, or under both conditions relative to the corresponding controls (Figure 3B). Several organs contained substantial proportions of Recspecific genes that were not differentially expressed immediately after stress. Incomplete return toward the control state therefore did not simply reflect persistence of the initial response; it also involved delayed or recovery-associated transcriptional programs.

Because the untreated transcriptomic states of young and aged organs were already different, having distinct DEG sets in the young and aged cohorts does not rule out the possibility that sleep deprivation is driving both cohorts to a common transcriptomic state where it can be best tolerated. We therefore used the difference between young and aged controls as a reference and asked whether the young–aged gap narrowed under sleep deprivation and recovery (Figure 3C). The brainstem and testis showed marked age-group convergence during recovery, with the young–aged difference decreasing by approximately 50% relative to that between the Ctrl groups. This result demonstrated that while some organs converged toward a common transcriptomic state in the face of sleep deprivation, others adopted a state that was highly age specific.

Finally, we evaluated transcriptomic displacement relative to the Young Ctrl and Aged Ctrl reference states separately (Figure 3D). The adrenal gland in young mice, the hypothalamus in both age groups, and the testis in aged mice moved away from both reference states, indicating changes extending beyond a simple young-to-aged axis. By contrast, the young testis and brain-stem shifted away from the Young Ctrl state and toward the Aged Ctrl state after sleep deprivation, and this displacement remained detectable after recovery. Consistent with the DEG-based analysis, the hypothalamus moved back toward its corresponding control state in both cohorts, whereas comparable recovery-associated changes were not observed in the aged heart and liver.

Together, these results show that the recovery period was not simply defined as the reversal of the initial stress response. Depending on the organ and age, the recovery period encompassed transcriptional rebound, persistence of stress-induced changes and delayed responses. In some cases, these complex trajectories can lead to convergence of young and aged transcriptomes toward a shared region of state space.

### Shape analysis and neural ODE modeling reveal distinct recovery landscapes and a separatrix

To quantify heterogeneous gene-expression trajectories, we defined a shape parameter:

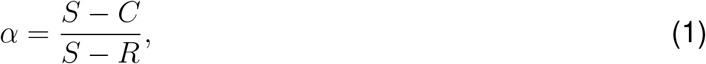

where *S, C* and *R* are the mean expression level of the Str, Ctrl and Rec group respectively. As its name suggests, *α* informs us about the shape of the expression dynamics (Figure 4A). For example, a gene with *α* = 1 indicates that it completely recovers from sleep deprivation (Figure 4A). As defined by the original equation, the shape parameter describes the expression dynamics of a gene (Figure 4A). Its distribution, therefore, summarizes whether genes returned toward control, remained displaced, or continued to change during recovery.

**Figure 4.**
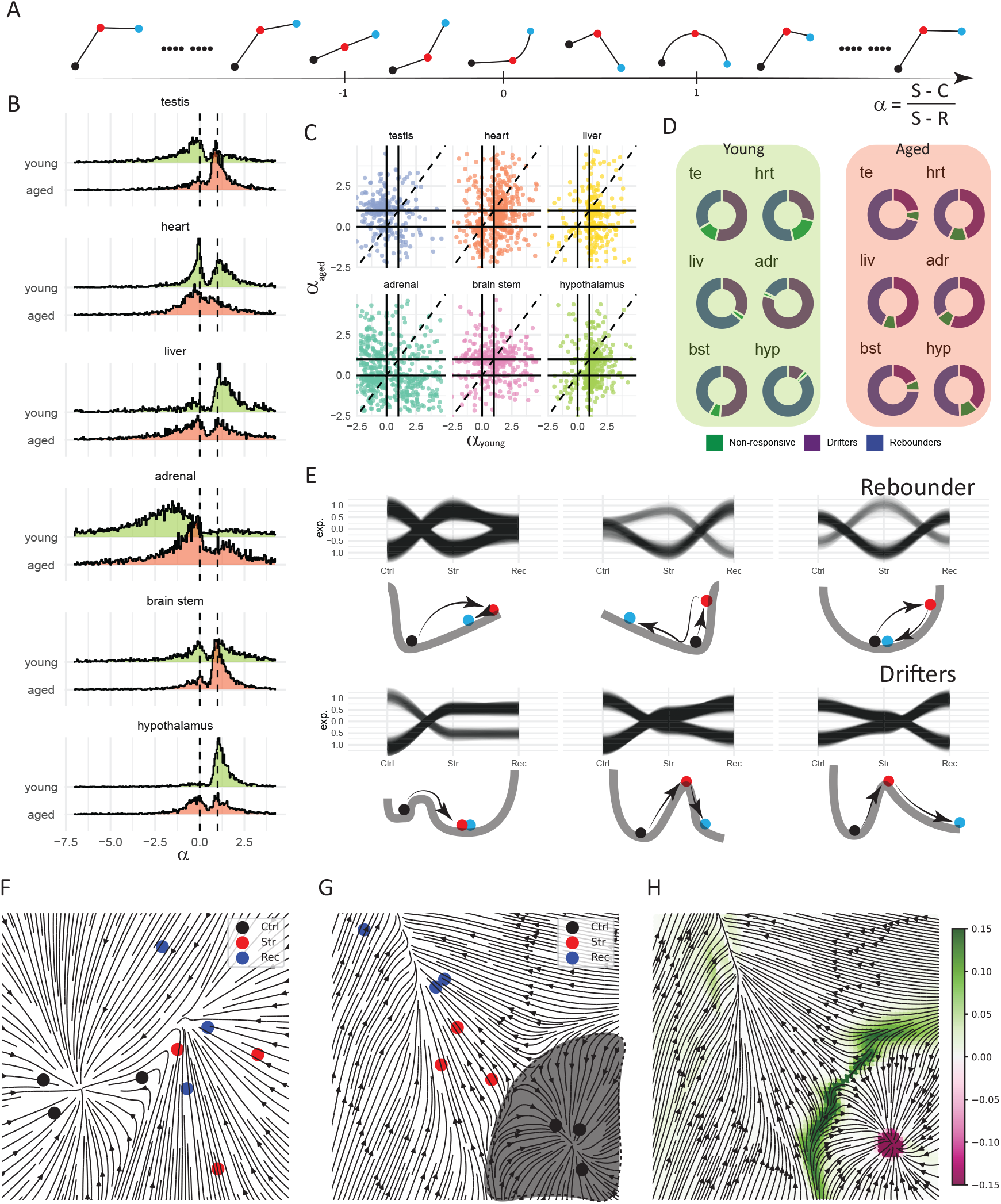
Shape analysis and neural ODE modeling identify distinct recovery regimes and a separatrix. (A)Definition of the shape parameter and schematic expression trajectories corresponding to different parameter values. The shape parameter was calculated using the mean expression levels of the stress (Str), control (Ctrl), and recovery (Rec) groups, as defined by the original equation. (B)Distributions of the shape parameter across genes in each organ from young and aged mice. Dashed lines indicate the parameter values corresponding to non-responsive genes and perfect recovery, as defined in (A). (C)Relationship between shape parameters calculated for the same sleep deprivation-responsive genes in young and aged mice across organs. (D)Fractions of rebounders, drifters, and non-responsive genes in each organ from young (left) and aged (right) mice. Categories were assigned according to the shape-parameter criteria defined in (A). (E)Representative expression trajectories of rebounders and drifters in the young testis. One-dimensional landscape illustrations corresponding to each trajectory class are shown below. These illustrations represent conceptual projections of the underlying high-dimensional transcriptomic landscape. (F)Neural ordinary differential equation (neural ODE)-inferred vector field in the principal component analysis (PCA) space of the young liver. The reconstructed field contained a single stable fixed point within the modeled region. Arrows indicate the inferred direction of transcriptomic state transition. (G)Neural ODE-inferred vector field in the PCA space of the young testis. The shaded region denotes the inferred basin from which trajectories return toward the Ctrl state. (H)Finite-time Lyapunov exponent (FTLE) calculated from the vector field in (G). The region of high perturbation sensitivity delineates the inferred separatrix bounding the basin of return to the Ctrl state. Ctrl, untreated control; Str, 24-h sleep deprivation; Rec, 24-h sleep deprivation followed by 3-day recovery.

In young mice, the liver and hypothalamus showed prominent peaks at *α* = 1, indicating strong signs of recovery, whereas the adrenal gland and testis lacked comparable peaks at the same position (Figure 4B). The brainstem and heart also exhibited peaks corresponding to continuous drift during the recovery period, consistent with the Rec-specific programs identified in Figure 3B. These patterns suggested delayed or recovery-associated responses but did not establish whether they originated locally or through interactions among organs. In aged mice, recovery-associated peaks remained detectable in the hypothalamus and liver but were reduced relative to those in young mice. Conversely, the aged adrenal gland and testis displayed more prominent recovery-associated peaks than their young counterparts. Aging therefore altered the distribution of gene-level recovery trajectories but did not uniformly impair transcriptional return across organs. For genes responsive to sleep deprivation in both age groups, shape parameters showed little correspondence between young and aged mice (Figure 4C). Thus, even when the same genes responded in both cohorts, their subsequent recovery trajectories were not necessarily preserved with age.

We classified the observed trajectories as rebounders, drifters, or non-responsive genes according to the original shape-parameter criteria (Figures 4D and 4E). Rebounders moved away from control after stress and subsequently returned toward it, whereas drifters remained displaced or continued to change during recovery. The proportions of these classes differed markedly across organs and ages. The one-dimensional illustrations in Figure 4E provide conceptual projections of these trajectory classes. Rebounders are represented by trajectories that return toward the initial state, whereas drifters move toward an alternative state. These projections should not be interpreted as direct evidence that an individual gene independently possesses one or multiple stable states. Functional enrichment analysis identified metabolism as a recurrent feature of rebounders in young mice. Carboxylic acid biosynthesis and organic acid biosynthesis were enriched in all six organs, and several additional metabolic processes were enriched in five of six organs. In aged mice, rebounders were associated with more heterogeneous functions, including chromatin remodeling, translation, metabolism, muscle contraction, and immune responses, with no term shared across all organs. Drifters were frequently enriched for muscle-related processes in both age groups. These findings identified distinct functional programs associated with transcriptional rebound and continued displacement.

Although the shape parameter summarized individual-gene trajectories, it could not reveal the organization of the multigene transcriptomic state space. We therefore performed principal component analysis separately for each organ and age cohort and used a neural ordinary differential equation (neural ODE) to infer vector fields describing organ-level state transitions during stress and recovery. As originally specified, the Ctrl state was defined as a stable fixed point, and points near the Str state were constrained to move toward the Rec state under the inferred flow. The reconstructed vector fields therefore represent data-constrained models of plausible state transitions rather than direct longitudinal measurements of individual animals. The reconstructed vector fields revealed two general stability regimes within the modeled regions. In the young liver, the field contained a single stable fixed point, with inferred trajectories directed toward the Ctrl state (Figure 4F). This organization was consistent with the prominent population of rebounders identified by the shape-parameter analysis. In contrast, the young testis would only return to the Ctrl state under weak perturbation (Figure 4G). The model identified a bounded region surrounding the control state within which the system can recover. If a perturbation pushed the system outside the identified bound, it would no longer return. Consistent with our visual inspection, finite time Lyapunov exponent (FTLE) of the state space revealed a region highly sensitive to perturbation, marking the boundary that determines whether the perturbed system can recover (Figure 4H). We refer to this inferred boundary as the separatrix. Within the reconstructed model, trajectories remaining inside the separatrix could return toward the Ctrl state, whereas trajectories crossing it would move away. Together, the shape-parameter and neural ODE analyses connected gene-level recovery behaviors to the inferred geometry of the transcriptomic landscape. They distinguished systems that are stable from those that are not, identifying a separatrix defining the boundary of regions inside which the perturbed system can return toward the control.

## DISCUSSION

Aging studies have generated increasingly detailed molecular atlases of young and aged tissues, but static comparisons provide limited information about the stability of these states^3–5^. Here, we used transient sleep deprivation followed by recovery to examine aging as a change not only in the transcriptomic state but also in the stress-induced gene expression dynamics. By integrating differential-expression analysis, a gene-level shape parameter, and neural ordinary differential equation modeling^15^ we developed a framework relating the observed stress–recovery responses to inferred transcriptomic landscapes^10,11^.

A central finding was that the same stressor did not produce a uniform response across organs or ages. The number, identity, and functional composition of sleep deprivation-responsive genes differed substantially among tissues, and genes responding in both young and aged mice frequently followed different recovery trajectories. Aging therefore did not simply amplify or attenuate a conserved stress-response program. Instead, it altered the transcriptional programs recruited by individual organs and the path taken after removal of the challenge. This observation extends multi-tissue aging atlases by showing that different organs are not only characterized by their baseline age-associated expression but also by their dynamic responses to perturbation.

In our study, we noted that recovery is not equivalent to the reversal of the initial stress response. Some genes returned toward control expression, whereas others remained displaced or only started to respond during recovery. These delayed responses may represent compensatory processes, secondary consequences of the initial perturbation, or transitions toward alternative states. Their presence emphasizes that measurements obtained immediately after stress cannot predict the eventual state of an organ. Accordingly, biological resilience is better represented by a trajectory following perturbation than by a single baseline or endpoint measurement^6,7,26^.

The shape-parameter analysis provided a concise description of this heterogeneity by distinguishing rebounders from drifters. Metabolic functions were recurrently enriched among rebounders in young organs, whereas muscle-related processes were frequently associated with drifters, particularly in aged organs. These patterns suggest that different biological programs vary in their propensity to return toward baseline. However, functional enrichment does not establish that metabolic programs drive recovery or that muscle-related programs cause persistent displacement. These possibilities require direct experimental perturbation.

Neural ODE modeling extended the gene-level analysis by reconstructing vector fields in a reduced transcriptomic state space^15^. The inferred fields distinguished systems that are “locally” stable from those that are “globally” stable. In the young testis, finite-time Lyapunov exponent^27,28^ analysis revealed a separatrix bounding the basin from which trajectories returned toward control. However, we acknowledge that computing FTLE on an inferred transcriptomic vector field requires cautious interpretation.

This framework has potential implications for intervention design. Conventional strategies often focus on reversing individual molecular abnormalities or increasing treatment magnitude. Our model instead suggests that intervention outcomes may depend jointly on the initial state and the direction and magnitude of the induced change. A treatment that shifts a system without returning it to the appropriate basin may fail to restore the original state, even if a handful of selected molecular markers is normalized. Conversely, a sufficiently directed intervention might move the system back inside the separatrix, after which intrinsic dynamics would drive the system to fully recover. Insights derived from our model reconstructed transcriptomic landscape echoed recent reports that successful state control requires positioning a system within an appropriate region of attraction^29^. In summary, our experimental design transforms aging from a static portrait into a dynamical map — one that reveals not just how systems change, but whether they can still come home. If the position of the separatrix can be learned, then aging intervention becomes a problem of control: nudging different organ systems back across the separatrix for self-driven rejuvenation.

### Limitations of the study

Several limitations define the scope of these conclusions. Ctrl, Str, and Rec samples were obtained from different animals at only three experimental conditions rather than through continuous longitudinal measurements. Bulk RNA sequencing cannot distinguish changes within cells from changes in cellular composition. The study included male mice and a single stress paradigm, limiting generalization across sexes, stressors, species, and disease states. The inferred vector fields depended on dimensionality reduction, modeling assumptions, and imposed stability constraints; alternative vector fields may also be compatible with the sampled states. Most importantly, the separatrix was computationally inferred and was not tested by experimentally applying perturbations on opposing sides of the predicted boundary. Future studies incorporating additional recovery time points, single-cell or spatial measurements, independent perturbations, and prospective testing of model-predicted trajectories will be required to determine whether the inferred boundaries represent general and controllable features of biological resilience.

## RESOURCE AVAILABILITY

### Lead contact

Further information and requests for resources should be directed to and will be fulfilled by the lead contact, Lei Li.

### Materials availability

This study did not generate new unique reagents.

### Data and code availability

RNA sequencing data generated in this study have been deposited in the Genome Sequence Archive with accession number CRA049304. Code generated in the analysis has been uploaded to https://github.com/bxxu/aging_landscape_stability. Any additional information required to reanalyze the data reported in this paper is available from the lead contact upon reasonable request.

## ACKNOWLEDGMENTS

This work was supported by the Shenzhen Medical Research Fund (SMRF-B2302013), the National Natural Science Foundation of China (NSFC32571311, NSFC32171154).

## AUTHOR CONTRIBUTIONS

Conceptualization, L.L. and X.B.; Methodology, L.L., X.B., and J.S.; Software, X.B.; Formal Analysis, X.B. and L.L.; Investigation, L.L. and J.S.; Data Curation, J.S., L.Z., D.C. and X.B.; Visualization, L.L. and X.B.; Writing – Original Draft, L.L. and X.B.; Writing – Review & Editing, L.L., X.B., J.S., D.C. and L.Z.; Supervision, L.L.; Project Administration, L.L.; Funding Acquisition, L.L. All authors reviewed and approved the final manuscript.

## DECLARATION OF INTERESTS

The authors declare no competing interests.

## STAR METHODS

### Experimental model and study participant details

#### Animals

All animal procedures were approved by the Institutional Animal Care and Use Committee of the Shenzhen Institute of Advanced Technology, Chinese Academy of Sciences. Animal experiments were conducted in accordance with applicable Chinese regulations and institutional guidelines and are reported following the ARRIVE guidelines 2.0.

Male wild-type C57BL/6J mice were used in this study (Guangdong Vital River Laboratory Animal Technology Co., Ltd., China). Young mice were 10–11 weeks old while aged mice were 24.5–30.5 months old at sample collection. Mice were group-housed in a specific pathogen-free facility under a 12-h light/12-h dark cycle, with food and water available *ad libitum*. Young and aged mice were randomly assigned to three experimental conditions: untreated control (Ctrl), 24-h sleep deprivation (Str), or 24-h sleep deprivation followed by 72-h recovery (Rec), with 3–5 mice initially assigned to each group. Following tissue collection, RNA samples were assessed for quantity, purity, and integrity before library construction. Samples that showed RNA degradation or otherwise failed the prespecified RNA-quality criteria were excluded before sequencing. The final numbers of mice represented in the transcriptomic dataset were Young Ctrl, n = 3; Young Str, n = 3; Young Rec, n = 3; Aged Ctrl, n = 3; Aged Str, n = 3; and Aged Rec, n = 2.

### Method details

#### Sleep deprivation and recovery

Sleep deprivation was performed using an automated rotating-bar apparatus (SD1900, Laiyue Biotech, China) operated at 3.5 revolutions per minute. The rotating bar moved continuously along the bottom of the chamber to prevent sustained sleep while allowing the mice unrestricted access to food and water. Mice in the Str groups underwent sleep deprivation for 24 h and were euthanized immediately after the challenge. Mice in the Rec groups underwent the same 24-h sleep-deprivation procedure and were subsequently returned to their home cages for 72 h before tissue collection. Ctrl mice remained untreated in their home cages before tissue collection.

#### Tissue collection

Mice were deeply anesthetized with 3% isoflurane delivered through a nose cone (Ringpu Tianjin Bio-Pharmacy Co., Ltd., China, absence of toe pinch reflex). Cervical dislocation was subsequently performed as the method of euthanasia. The hypothalamus, brainstem, left testis, bilateral adrenal glands, heart, and liver were then collected. The bilateral adrenal glands were pooled to generate one adrenal sample per animal. The hypothalamus and brainstem were dissected according to anatomical landmarks. All tissues were immediately snap-frozen in liquid nitrogen and stored at − 80°C until RNA extraction and further sequencing analysis (LC Bio Technology Co., Ltd., Hangzhou, China).

#### RNA extraction and quality control

Total RNA was extracted from frozen tissues using TRIzol reagent (Thermo Fisher Scientific, catalog no. 15596018) according to the manufacturer’s instructions. RNA quantity, purity, and integrity were assessed using an Agilent 2100 Bioanalyzer with the RNA 6000 Nano LabChip Kit (Agilent Technologies, catalog no. 5067-1511). Only RNA samples with an RNA integrity number greater than 7.0 were used for library construction. Samples showing evidence of RNA degradation did not meet the prespecified quality-control criterion and were excluded before library construction.

#### RNA library preparation and sequencing

Polyadenylated RNA was isolated from 5 *μ*g total RNA using Dynabeads Oligo(dT) (Thermo Fisher Scientific) through two rounds of purification. Purified messenger RNA was fragmented at 94°C for 5–7 min using the Magnesium RNA Fragmentation Module (New England Biolabs, catalog no. E6150). First-strand complementary DNA was synthesized using SuperScript II Reverse Transcriptase (Invitrogen, catalog no. 1896649). Uracil-containing second-strand complementary DNA was synthesized using *Escherichia coli* DNA polymerase I (New England Biolabs, catalog no. M0209), RNase H (New England Biolabs, catalog no. M0297), and deoxyuridine triphosphate solution (Thermo Fisher Scientific, catalog no. R0133). Following end repair and A-tailing, dual-index adapters were ligated, and fragments were size-selected using AMPure XP beads. Uracil-containing second strands were digested using heat-labile uracil-DNA glycosylase (New England Biolabs, catalog no. M0280). Libraries were amplified by polymerase chain reaction using the following program: 95°C for 3 min; eight cycles of 98°C for 15 s, 60°C for 15 s, and 72°C for 30 s; and 72°C for 5 min. The final complementary DNA libraries had an average insert size of 300 *±* 50 bp. Libraries were sequenced as 150-bp paired-end reads on an Illumina NovaSeq 6000 platform by LC-Bio Technology Co., Ltd. (Hangzhou, China).

Adapter-containing and low-quality reads were removed using Cutadapt version 1.9. Reads containing poly(A) or poly(G) sequences, more than 5% undetermined nucleotides, or more than 20% bases with a Phred quality score of 20 or lower were excluded. The quality of the cleaned reads was assessed using FastQC version 0.11.9, including Q20, Q30, and guanine-cytosine content. Clean reads were aligned to the mouse reference genome GRCm38/mm10 using HISAT2 version 2.2.1. Transcript assembly and expression quantification were performed using StringTie version 2.1.6. Transcript assemblies from all samples were merged using gffcompare version 0.9.8. Expression abundance was calculated as fragments per kilobase of transcript per million mapped reads for visualization and trajectory analyses where specified.

#### Bioinformatic analysis

Differentially expressed genes were identified via the DESeq2^30^ package with default parameters. Over-representation analysis and gene set enrichment analysis were performed using the clusterProfiler^31,32^ package with the “compareCluster” and “gseGO” functions respectively with default parameters.

#### Computing the shape parameter and classifying genes

The shape parameter is defined as 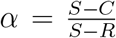, where *S, C* and *R* is the mean expression level of the Str, Ctrl and Rec group respectively. The shape parameter was calculated for a gene if it showed differential expression between the Ctrl and Str group, Ctrl and Rec group or both. The subsequent distribution of the shape parameter also consists of the same genes. To visualize genes with similar shape parameter, FPKM of each gene was first z-scored and fitted via loess regression using the ggplot2^33^ package in R. A gene is considered to be a rebounder if its shape parameter ranges from 0.2 to 10 or a drifter if its shape parameter has an absolute value greater than 10 or less than −0.2. Genes with *α* between −0.2 to 0.2 are considered to be non-responsive to sleep deprivation.

#### Characterizing local landscape using neural ODE

A neural ODE is defined by a neural network *f* with parameters *θ* which defines the motion on the phase space as 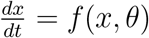. The network is trained using the backward adjoint dynamics. To reconstruct a landscape that agrees with our data, we make the assumption that the Ctrl state is stable. By initiating points, 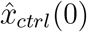, around the mean of the Ctrl state, *x*_*ctrl*_, we subjected 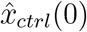 to the dynamics defined by the network for a total duration of *T*_*ctrl*_ and defined the control state stability loss as:

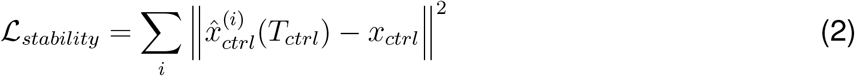

to enforce the stability of the Ctrl state. To enforce a flow from the Str state to the Rec state, we initiated points, 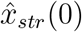, around the mean of the Str state. Similarly, we subjected initiated points to the dynamics defined by the network for a total duration of *T*_*str*_, and defined the recovery dynamics loss as:

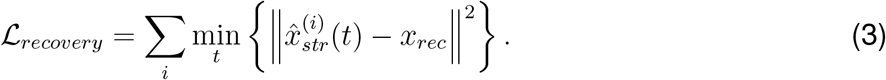

We employed a fully connected network with RELU activation using PyTorch^34^ and conducted training using the torchdiffeq^15^ package.

#### Computing the finite time Lyapunov exponent

Finite time Lyapunov exponent was computed using custom Python code. In short, we first initialize a grid of points on the phase space, and compute the flow map Jacobian, defined as:

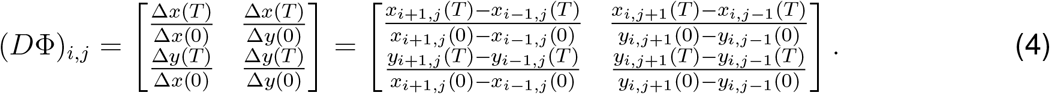

Next, we compute the Cauchy-Green deformation tensor:

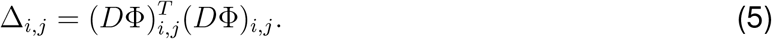

Lastly, the finite time Lyapunov exponent at (*x*_*i,j*_, *y*_*i,j*_) is defined as:

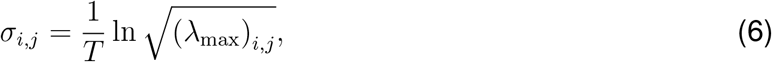

where (*λ*_max_)_*i,j*_ is the maximum eigenvalue of the Cauchy-Green deformation tensor.

#### Mathematical characterization of the stability of a fixed point

Given a system of differential equations in the form:

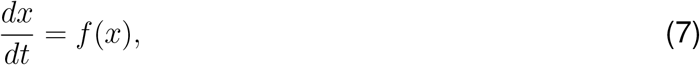

with steady states *x*^*\**^ such that *f* (*x*^*\**^) = 0. The stability of its steady states is probed through the prescription of an infinitesimal perturbation and study whether the perturbed system relaxes back to the steady state *x*^*\**^. Mathematically, the perturbation is written as:

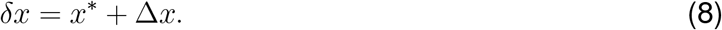

By definition, the dynamics of *δx* is governed by *f*(*•*), namely:

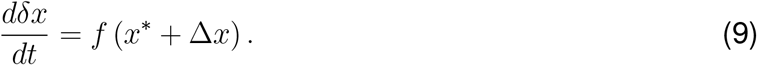

*f* (*x*^*\**^ + Δ*x*) can be expanded via Taylor expansion; dropping higher order terms, we obtain a linear system:

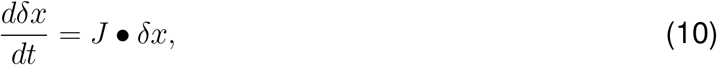

with solution:

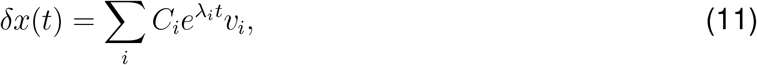

where *J, C*_*i*_, *λ*_*i*_, *v*_*i*_ denotes the Jacobian matrix (defined as 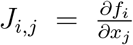), constant terms, eigenvalue and its associated eigenvector of *J* respectively. Here, we can tell easily that the system is unstable if a single eigenvalue of the Jacobian matrix is greater than 0.

#### Relationship between the gene expression landscape and noise

Noise in dynamical systems can be modeled as:

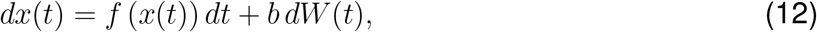

where *W* defines a Wiener process. By setting *W* (0) to zero, we would have *W* (*t*) ~ 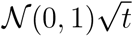 where *N* (0, 1) is Gaussian with zero mean and unit variance. To get the time evolution of *x*, one can integrate the above expression in a discrete manner:

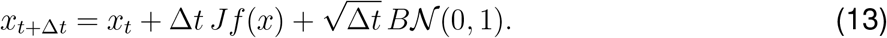

Here, *A* is the Jacobian matrix and *B* denotes the covariance matrix of the noise injected to the system. The reason why we have the Jacobian matrix here is that we are again interested in the dynamics near the fixed points of the system. From this expression, we can infer the covariance matrix *C*, defined as *C*_*i,j*_ = *E*(*x*_*i*_ *• x*_*j*_), where *E*(*•*) denotes expectation. Denoting the multivariate Gaussian *BN* (0, 1) as *ξ* and its covariance *D* = *BB*^*T*^, the time evolution of the covariance matrix is defined as:

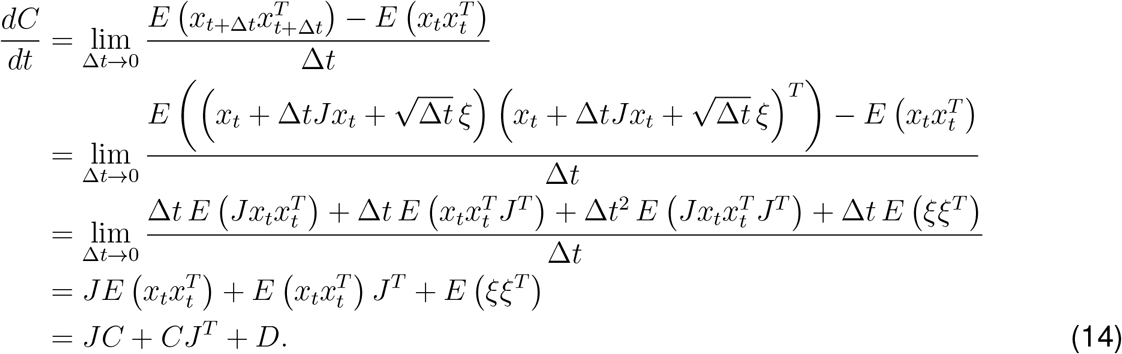

At steady state, we have the continuous time Lyapunov equation:

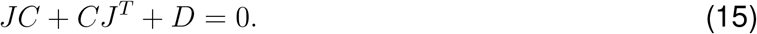

If we assume that the Jacobian matrix can be decomposed as:

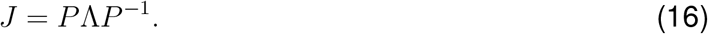

In this expression, *P* is a matrix whose columns 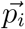 denotes the eigenvectors of the original matrix and Λ is a diagonal matrix of eigenvalues. Left-multiply the Lyapunov equation with *P* ^*−*1^ and right-multiply with (*P*^*T*^)^*−*1^, we get:

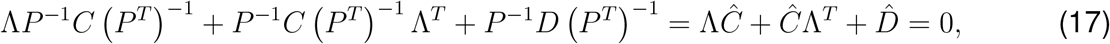

where *Ĉ* = *P* ^*−*1^*C (P*^*T*^) ^*−*1^ and 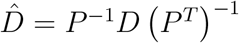.

The above equation can be solved in an element-wise fashion:

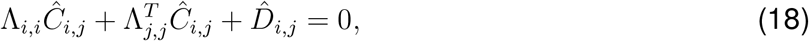

from which we get:

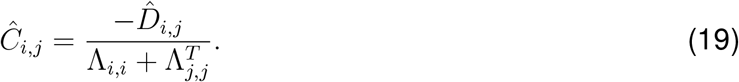

Transforming *Ĉ*_*i,j*_ back to the covariance matrix *C*_*i,j*_, we see that as the largest eigenvalue of *A* approaches 0, assuming Λ_1,1_ is the largest eigenvalue of the Jacobian matrix,

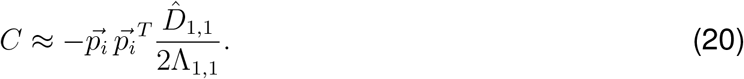

## Notes

### Competing Interest Statement

The authors have declared no competing interest.

